# Grain Protein Content QTLs Identified in a Durum × Wild Emmer Wheat Mapping Population Tested in Five Environments

**DOI:** 10.1101/601773

**Authors:** Andrii Fatiukha, Itamar Lupo, Gabriel Lidzbarsky, Valentina Klymiuk, Abraham B. Korol, Curtis Pozniak, Tzion Fahima, Tamar Krugman

## Abstract

**Abstract:** Wild emmer wheat (*Triticum turgidum* ssp. *dicoccoides*, WEW) was shown to exhibit high grain protein content (GPC) and therefore, possess a great potential for improvement of cultivated wheat nutritional value. A recombinant inbred line (RIL) population derived from a cross between *T. durum* var. Svevo and WEW acc. Y12-3 was used for construction of a high-density genetic map and genetic dissection of GPC. Genotyping of 208 F_6_ RILs with 15K wheat SNP array yielded 4,166 polymorphic SNP markers, of which 1,510 were designated as skeleton markers. A total map length of 2,169 cM was obtained with an average distance of 1.5 cM between SNPs. A total of 12 GPC QTLs with LOD score range of 2.7-35.9, and PEV of 2.6-26.6% were identified under five environments. Major QTLs with favorable alleles from WEW were identified on chromosomes 4BS, 5AS, 6BS and 7BL. The QTL region on 6BS coincided with the physical position of the previously cloned QTL, *Gpc-B1*. Comparisons of the physical intervals of the GPC QTLs described here with the results previously reported in other durum×WEW RIL population led to the identification of four common and two homoeologous QTLs. Exploration of the large genetic variation within WEW accessions is a precondition for discovery of exotic beneficial alleles, as we have demonstrated here, by the identification of seven novel GPC QTLs. Therefore, our research emphasizes the importance of GPC QTL dissection in diverse WEW accessions as a source of novel alleles for improvement of GPC in cultivated wheat.

**Key message:** Genetic dissection of GPC in tetraploid durum × WEW RIL population, based on high-density SNP genetic map, revealed 12 QTLs, with favorable WEW allele for 11 QTLs.

## Introduction

Common wheat (*Triticum aestivum* L.) is one of the most important crops with world production of more than 770 million tons harvested over 220 million hectares of land area in 2017 (OECD/FAO 2018). Wheat is a rich source of carbohydrates, however wheat grains contain rather moderate amount of proteins, which is ranging from 9 to 15% only (Shewry and Hey 2015). Grain protein content (GPC) shapes the diet nutritional value important for human health and partially determines the baking properties of bread wheat, as well as the pasta-making characteristics of durum wheat (*Triticum turgidum* L. subsp. *durum*) (Blanco et al. 2012). Proteins and carbohydrates are the main components composing grain weight; each has its own limiting environmental conditions, including (i) availability of utilized nitrogen for protein accumulation, and (ii) sufficient levels of water and sunlight required for CO2 fixation and carbohydrate synthesis. Environmental factors (biotic and abiotic) and their interactions with different genetic backgrounds (genotypes) were shown to affect significantly the total amount and composition of grain proteins (Triboï□Blondel et al. 2003). Drought is one of the major abiotic stresses, mainly affecting grain carbohydrate content (Balla et al. 2011), it also plays an important role as a major environmental factor influencing GPC (Flagella et al. 2010). Global climate change is intensifying the severity of drought around the word and, thus, requires breeding of crops for adaptation to this environmental stress (Asseng et al. 2014).

The evolutionary forces that exerted on natural populations of crop wild relative have shaped their genetic structure leading to their superior adaptation to various environments including those with water-limited conditions. Therefore, the exploitation of their rich genetic repertoire has an extensive potential for crop improvement (Henry and Nevo 2014; Krugman et al. 2018; Klymiuk et al. 2019b). Wild emmer wheat (WEW) (*Triticum turgidum* L. subsp. *dicoccoides*) is the tetraploid progenitor of both, tetraploid durum and hexaploid common wheats. WEW germplasm represents a valuable reservoir of genetic variation for drought resistance and GPC (Huang et al. 2016; Klymiuk et al. 2019a) that may serve as a promising source of favorable alleles for wheat breeding. One such WEW genotype is Y12-3, that was shown to be resistant to drought (Peleg et al. 2005) and confer high GPC and mineral content (Peleg et al. 2008). Transcriptomic and metabolomic profiles of this genotype were extensively studied under terminal drought stress (Krugman et al. 2010, 2011). Y12-3 was used for the development of the recombinant inbred line (RIL) population used in the current study.

*Gpc-B1* is a high GPC QTL, assigned to chromosome arm 6BS, with GPC increasing allele originated from WEW (Joppa et al. 1997). It was shown to be associated also with increased grain zinc/iron content and earlier leaf senescence, and to be controlled by the transcription factor *TtNAM-B1* (Uauy et al. 2006). In tetraploid wheat, two additional copies were found on chromosome arms 6AS (*TtNAM-A1*, an orthologous copy), and chromosome arm 2BS (*TtNAM-B2*, paralogous copy, that is 91% identical to *TtNAM-B1* at the DNA level), while hexaploid wheat genome contains four *TaNAM* copies (*TaNAM-A1, D1, B2*, and *D2*; Uauy et al. 2006). WEW allele of *Gpc-B1* showed positive effects on GPC and other quality traits, and minor impacts on yield related traits, following its introgression into various wheat backgrounds (Brevis and Dubcovsky 2010; Tabbita et al. 2017).

High-throughput single nucleotide polymorphism (SNP) genotyping is a powerful tool that can assist in improving wheat genetic maps and construction of SNP-based consensus maps of tetraploid (Maccaferri et al. 2015) and hexaploid wheat (Wang et al. 2014). SNP-based arrays developed for bread wheat (e.g. the 90K and 15K arrays) were proved to be highly efficient also for construction of genetic maps based on WEW as one of the parents, including studies related to wheat chromosome evolution (Jorgensen et al. 2017), wheat domestication (Nave et al. 2016; Golan et al. 2018), QTL analysis of drought adaptation (Fatiukha et al. 2019a) and nutrient quality of wheat grains (Fatiukha et al. 2019b). In addition, high-density genetic map of durum Svevo × WEW Zavitan (S×Z) RIL population (Avni et al. 2014a) served as a base for anchoring of WEW genome assembly scaffolds (Avni et al. 2017). QTL analysis is a suitable approach for genetic dissection of complex traits such as GPC. The use of SNP-based maps for QTL analysis allows to narrow down the intervals of the detected QTLs compared to previous SSR-based maps (Fatiukha et al. 2019a) and to compare results obtained from different populations, and can serve as a powerful tool for marker-assisted selection (MAS) of QTLs during crop breeding programs.

In the current study the RIL population (S×Y) derived from a cross between durum wheat Svevo and WEW Y12-3 (S×Y) and genotyped with 15K SNP array, was used for construction of high-density genetic map and further QTL analysis of GPC. Phenotypic data was obtained from a total of five environments (three locations two of them with two contrasting water regimes). QTL analysis revealed 12 GPC QTLs, for which 11 favorable QTL alleles were contributed by WEW. We further used the whole genome assembly of WEW (Avni et al. 2017) for localization of candidate genes (CGs) associated with GPC, residing within the obtained QTL intervals. In addition, the physical positions of mapped SNPs and lists of CGs from the QTL intervals were used for identification of common and homoeologous CGs between the two durum × WEW RIL populations.

## Materials and methods

### Plant material, growth conditions, and phenotyping

A RIL population (S×Y) of 208 F_6_ lines was derived from the cross between an elite durum cultivar Svevo (Arduini et al. 2006; Maccaferri et al. 2008) and WEW accession Y12-3 (Peleg et al. 2005; Krugman et al. 2010) using the single seed descend approach. The RIL population was tested under a total of five environments in three sites over 2 years across Israel: i) an open field at Ein-Tamar, during 2013-2014 winter season, (30°56”N, 35°22”E), on calcite cierozems soil; ii) an open field at Kimaron during 2014-2015 winter season (32°29”N, 35°30”E), on light brown alluvian and clay-like silt sierozems; iii) water-sheltered green-house at Sharona during 2014-2015 winter season, (32°43”N, 35°27”E), on rendzina soil. In Ein Tamar (ET) annual rainfall was only 66 mm, therefore additional 430 mm were applied by drip irrigation (in total 496 mm). In Kimaron two water regimes were applied: well-watered (230 of rain, supplemented with 622 mm by drip irrigation, a total of 852 mm; K_WW) and water-limited (230 of rain, supplemented with 208 mm by drip irrigation, a total of 438 mm; K_WL). In Sharona taking into account high level of ground water two water regimes were applied: well-watered (supplemented with 448 mm by drip irrigation; S_WW) and water-limited (supplemented with 144 mm by drip irrigation; S_WL). Each of the 208 RILs and parental lines were represented by four plants in each plot (15×20 cm) with three replicates, considered as an experimental unit with randomize block design. Grain protein concentrations were measured using Leco N x 5.7 (Leco Corp., St. Joseph. MI) (Am. Assoc. Cereal Chem. Method 46–30).

### DNA extraction and SNP genotyping

The fresh leaf tissues of the parental genotypes (Svevo and Y12-3) and each of the 208 F_6_ RILs were used for DNA extraction following a standard CTAB protocol (Doyle 1991). DNA was normalized to 50 ng/μl. SNP genotyping was performed using the Illumina Infinium 15K Wheat platform, developed by TraitGenetics, Gatersleben, Germany (Muqaddasi et al. 2017), consisting of 12,905 SNPs selected from the wheat 90K array (Wang et al. 2014).

### Statistical analysis of phenotypic data

Statistical analysis that included ANOVA and testing for normal distribution was performed using the BioVinci software (BioTuring, San Diego, CA, USA).

### Construction of high-density genetic map

The genetic map was constructed using MultiPoint software, version «UltraDense» (http://www.multiqtl.com) (Ronin et al. 2017). The best candidate skeleton markers representing groups of co-segregating markers with size of ≥2 were selected using the function “bound together” after filtering for missing data (removing markers with more than 10% missing data points) and large segregation distortion (χ^2^ > 38). The threshold of recombination fraction (RF=0.2) was applied for clustering of candidate markers into linkage groups (LG). Marker ordering and testing of the local map stability and monotonicity were performed for each LG (Mester et al. 2003; Korol et al. 2009). The final number of LGs was reduced to 14, in accordance with the haploid number of chromosomes of tetraploid wheat, by merging the LGs with minimum pairwise RF values. The correspondence of the mapped markers with those on the consensus maps of hexaploid (Wang et al. 2014) and tetraploid wheat (Maccaferri et al. 2015) were used for orientation of each LG in relation to the short (S) and long (L) chromosome arms.

### QTL analysis

QTL analysis was performed using the general interval mapping (IM) procedure of MultiQTL software package (http://www.multiqtl.com). Single-QTL and two-linked-QTL models (Korol et al. 2009) were employed for screening of genetic linkage for GPC in each environment separately. Multi-environment analysis (MEA) was performed by joint analysis of trait values scored in five environments. After independent analysis for each chromosome, multiple interval mapping (MIM) was used to reduce the residual variation for each QTL under consideration, while taking into account QTLs that reside on other chromosomes (Kao et al. 1999). The significance of the detected QTL effects was tested using 10000 permutation runs. Significant models were further analyzed by 10000 bootstrap runs to estimate the standard error of the QTL effect.

### Identification of the physical intervals of QTLs and CGs

The physical positions of SNP markers were obtained by BLAST search of probe sequences (Wang et al. 2014) against the whole-genome assembly of WEW accession ‘Zavitan’ (Avni et al. 2017). Annotated gene models of the ‘Zavitan’ genome assembly (Avni et al. 2017) were used for identification of genes residing within QTL intervals (1.5 LOD support interval of QTL).

## Results

### High-density genetic map

Genotyping of the S×Y RIL population yielded 4,537 segregating SNP markers, out of these, 4,164 SNPs represented 1,510 unique loci (skeleton markers) were clustered into 14 LGs (Fig. 1). The constructed genetic map covered 2,168.6 cM with approximately equal length for the A (1,091.7 cM) and B genomes (1,076.9 cM) (Table 1, Table S1). The length of the individual chromosome maps ranged, from 114.2 cM (1A) to 188.5 (5A) and the number of skeletal markers ranged from 78 (4B) to 148 (2B). In total 117 (4.04%) non-recombinant chromosomes were observed among 208×14=2,912 RIL × chromosome combinations (Table 2), of which 50 were homogeneous for the WEW parental alleles and 67 for the durum parental alleles. The highest numbers of non-recombinant chromosomes were found for chromosomes 1B (28) and the lowest for 7A (2). Genome A showed significant (*P*≤.05) higher number of non-recombinant chromosomes compare to the B genome. A significant negative correlation, R=-0.78 (*P*<0.05), was found between the proportion of non-recombinant chromosomes and the length of the individual chromosome maps (Fig. S1). A total of 505 (33%) skeletal loci showed significant (*P*≤0.05) segregation distortion (Fig. S2), 71 loci in favor of Y12-3 and 434 loci for Svevo. Moreover, an extremely higher number of these loci was from the B genome than the A genome (351 vs 154). The highest number of distorted markers (87) was found for chr. 2B, while chr. 4B did not show any distorted marker. More than 93% of the markers (3882 out of 4164) mapped in the S×Y population were anchored to the reference genome of tetraploid WEW (Avni et al., 2017) (Table S1). High collinearity with average rank correlation coefficient of 0.98 was observed between the genetic and the physical positions of the mapped SNPs (Fig. S3).

**Figure 1.**
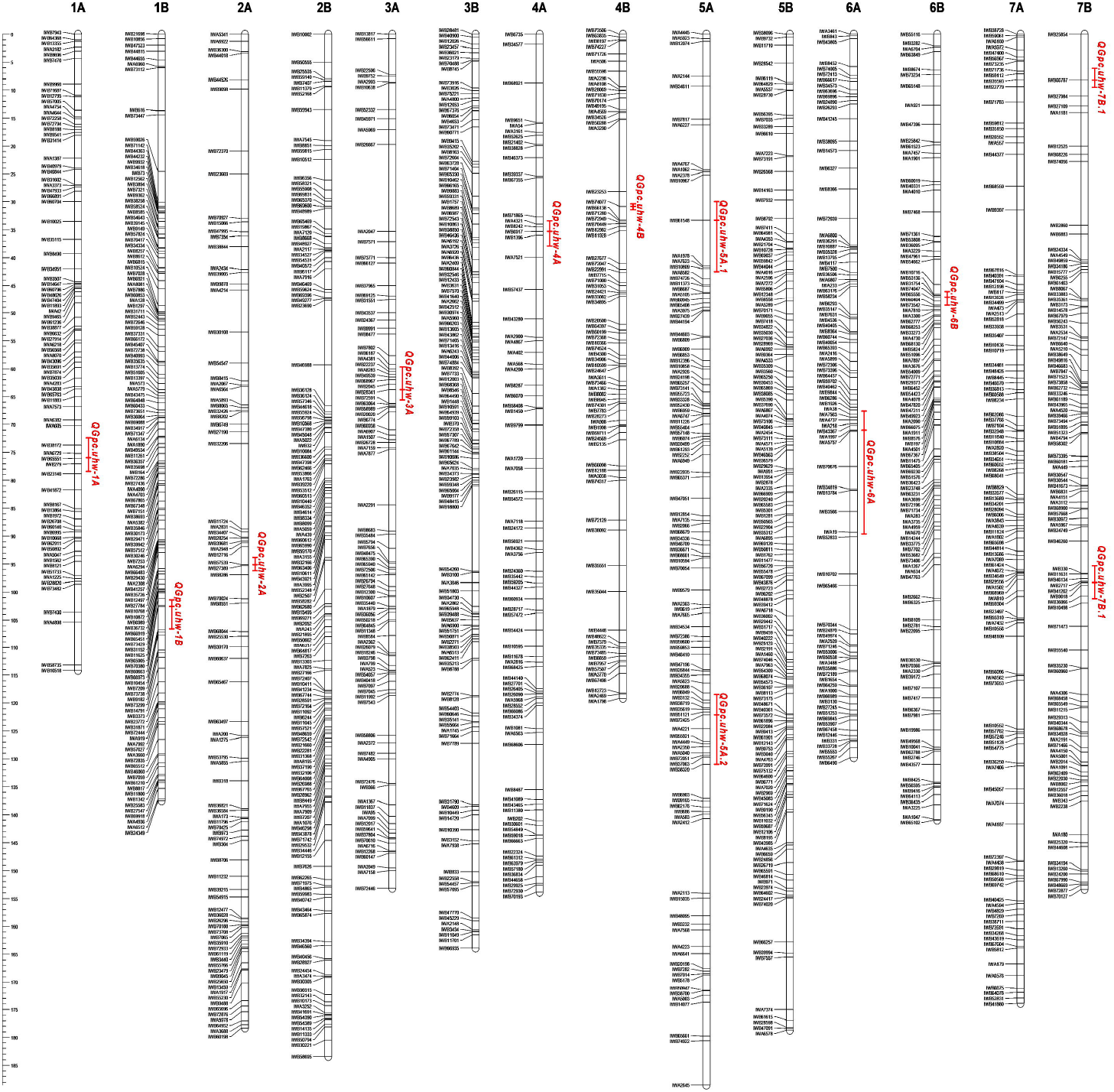
High-density genetic map of S×Y RIL population and 1.5 LOD support intervals of GPC QTLs detected under five environments.

**Table 1.** Summary of the genetic map constructed based on S×Y RIL population

| Linkage group | Total markers | Skeletal markers | Co-segregated markers | Length (cM) | Average interval (cM)/locus |
| --- | --- | --- | --- | --- | --- |
| 1A | 230 | 81 | 149 | 114.2 | 1.43 |
| 1B | 381 | 135 | 246 | 137.5 | 1.03 |
| 2A | 240 | 88 | 152 | 178.3 | 2.05 |
| 2B | 423 | 148 | 275 | 183.4 | 1.25 |
| 3A | 254 | 94 | 160 | 153.2 | 1.65 |
| 3B | 365 | 130 | 235 | 163.9 | 1.27 |
| 4A | 162 | 80 | 82 | 153.9 | 1.95 |
| 4B | 169 | 78 | 91 | 119.2 | 1.55 |
| 5A | 287 | 113 | 174 | 188.5 | 1.68 |
| 5B | 408 | 149 | 259 | 178.7 | 1.21 |
| 6A | 310 | 90 | 220 | 129.8 | 1.46 |
| 6B | 358 | 106 | 252 | 141.0 | 1.34 |
| 7A | 295 | 114 | 181 | 173.9 | 1.54 |
| 7B | 284 | 104 | 180 | 153.4 | 1.49 |
| Group 1 | 611 | 216 | 395 | 251.7 | 1.23 |
| Group 2 | 663 | 236 | 427 | 361.6 | 1.65 |
| Group 3 | 619 | 224 | 395 | 317.1 | 1.46 |
| Group 4 | 331 | 158 | 173 | 273.1 | 1.75 |
| Group 5 | 695 | 262 | 433 | 367.1 | 1.44 |
| Group 6 | 668 | 196 | 472 | 270.8 | 1.40 |
| Group 7 | 579 | 218 | 361 | 327.3 | 1.51 |
| A genome | 1778 | 660 | 1118 | 1091.7 | 1.68 |
| B genome | 2388 | 850 | 1538 | 1076.9 | 1.30 |
| **Total** | **4166** | **1510** | **2656** | **2168.6** | **1.49** |

**Table 2.** Number of RILs with parental (non-recombinant) chromosomes in S×Y RIL population

| Chromosome | Svevo | Y12-3 | Total | % RILs |
| --- | --- | --- | --- | --- |
| 1A | 14 | 14 | 28 | 13.53 |
| 1B | 8 | 2 | 10 | 4.83 |
| 2A | 5 | 1 | 6 | 2.90 |
| 2B | 2 | 2 | 4 | 1.93 |
| 3A | 4 | 4 | 8 | 3.86 |
| 3B | 4 | 2 | 6 | 2.90 |
| 4A | 5 | 6 | 11 | 5.31 |
| 4B | 7 | 8 | 15 | 7.25 |
| 5A | 3 | 1 | 4 | 1.93 |
| 5B | 2 | 1 | 3 | 1.45 |
| 6A | 8 | 5 | 13 | 6.28 |
| 6B | 2 | 1 | 3 | 1.45 |
| 7A | 1 | 1 | 2 | 0.97 |
| 7B | 2 | 2 | 4 | 1.93 |
| Genome A | 40 | 32 | 72 | 4.97 |
| Genome B | 27 | 18 | 45 | 3.11 |
| Total | 67 | 50 | 117 | 4.04 |

### GPC of the RIL population

The wild parent Y12-3 showed on average ~42 % higher GPC than Svevo (21.4 vs. 15.1), under all five environments (ET, K_WL, K_WW, S_WL, and S_WW) (Table 3), while the RILs exhibited a wide range (13.9—29.2) under the five environments over 2 years (Fig. 2, Table 3). GPC of parental lines and RILs was lower under the green-house conditions (S_WL and S_WW) compared to the field conditions (ET, K_WL, and K_WW). The highest GPC values for the RILs was obtained under ET (21.8), while the lowest GPC was detected under S_WL (17.9) (Fig. 1, Table 3). Transgressive segregation of GPC was obtained for the RIL population only under K_WL environment, whereas under other four environments GPC of Svevo was lower than that of RILs (Fig. S4). GPC values across the five environments approximately fits normal distribution under all environments (Fig. S4). Analyses of variance (ANOVA) showed highly significant effects (*P*≤0.001) for genotype, irrigation regimes, and environments for GPC (Table 3), while the genotype × environment interactions were not significant. Heritability (h^2^) calculated across environments was relatively high (0.77) (Table 3).

**Figure 2.**
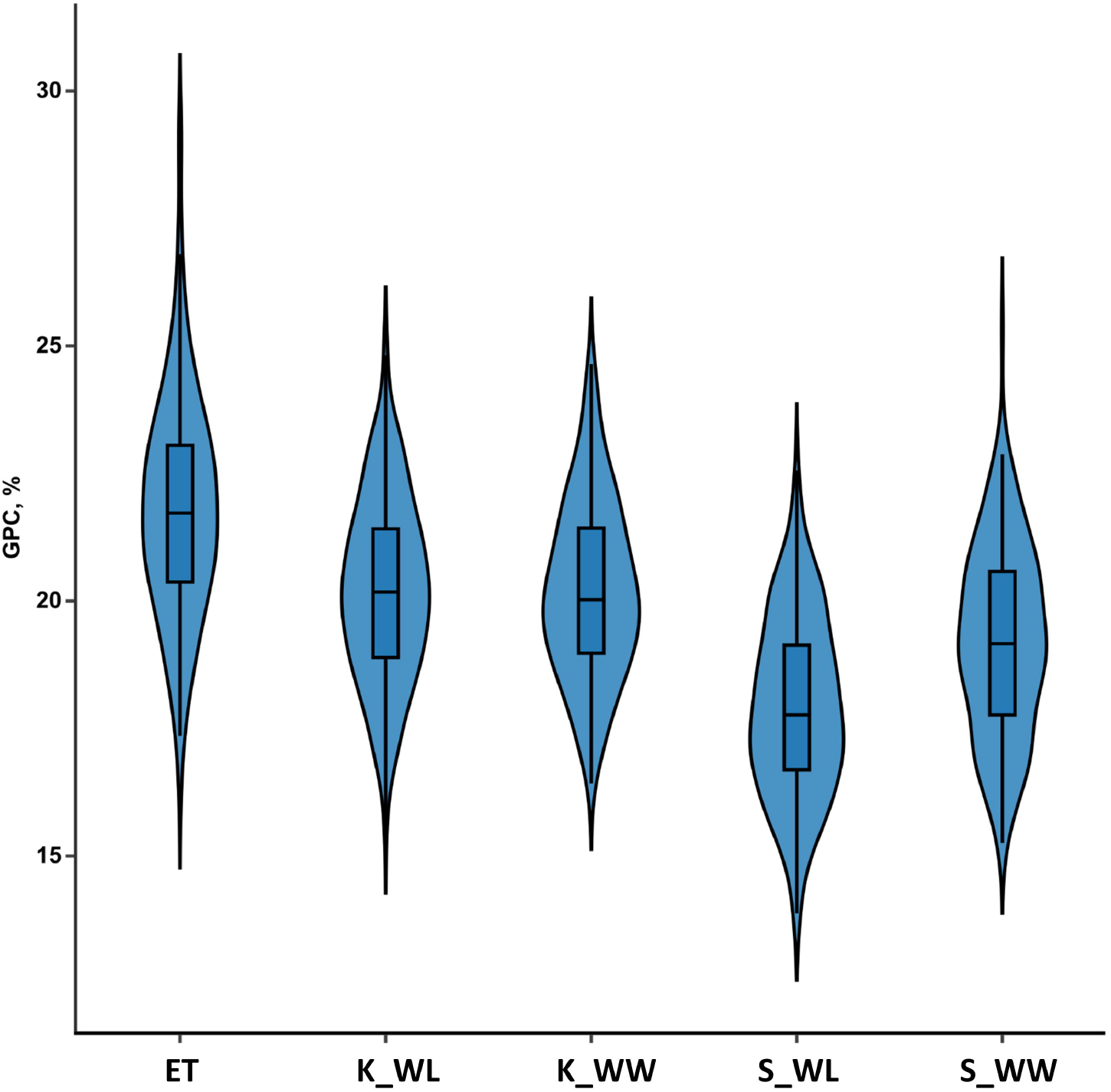
Distribution of GPC in S×Y RIL population. The violin plots show the distribution of GPC in wheat grains measured across five environments: open fields at Ein-Tamar and Kimaron (ET, K_WL, and K_WW) and green-house at Sharona (S_WL and S_WW). Suffixes WL and WW designate well-watered and water-limited conditions.

**Table 3.**
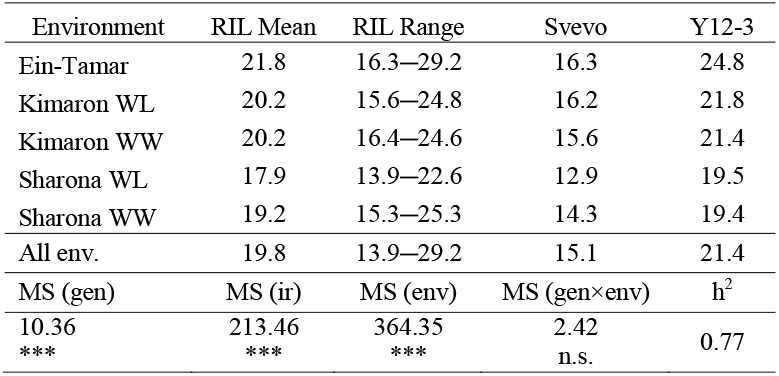
Analyses of variance (ANOVA), means and ranges for grain protein content (GPC) in S×Y RIL population under five environments.

| Environment | RIL Mean | RIL Range | Svevo | Y12-3 |
| --- | --- | --- | --- | --- |
| Ein-Tamar | 21.8 | 16.3—29.2 | 16.3 | 24.8 |
| Kimaron WL | 20.2 | 15.6—24.8 | 16.2 | 21.8 |
| Kimaron WW | 20.2 | 16.4—24.6 | 15.6 | 21.4 |
| Sharona WL | 17.9 | 13.9—22.6 | 12.9 | 19.5 |
| Sharona WW | 19.2 | 15.3—25.3 | 14.3 | 19.4 |
| All env. | 19.8 | 13.9—29.2 | 15.1 | 21.4 |
| MS (gen) | MS (ir) | MS (env) | MS (gen×env) | h<sup>2</sup> |
| 10.36 | 213.46 | 364.35 | 2.42 | 0.77 |
| *** | *** | *** | n.s. |  |

### Identification of GPC QTLs

A total of 12 significant GPC QTLs were detected in the S×Y RIL population with LOD scores range of 3.6-27.8 and percent of explained variation (PEV) range of 0.6-24.4% (Table 4). Y12-3 allele contributed to increasing of GPC at 11 QTLs, while Svevo allele contributed only to one (*QGpc.uhw-1B*). Most of the QTLs had significant effects under all environments, although three QTLs (*QGpc.uhw-5A.2, QGpc.uhw-6A*, and *QGpc.uhw-7B.1*) were significant only under the greenhouse conditions. *QGpc.uhw-4B, QGpc.uhw-5A.1, QGpc.uhw-6B*, and *QGpc.uhw-7B.2* seems to be major GPC QTLs with the highest LOD scores and relatively high and stable PEV for most of the environments. Interestingly, seven of the detected QTLs resided on chromosomes of genome A, and only five on genome B (Table 4).

**Table 4.**
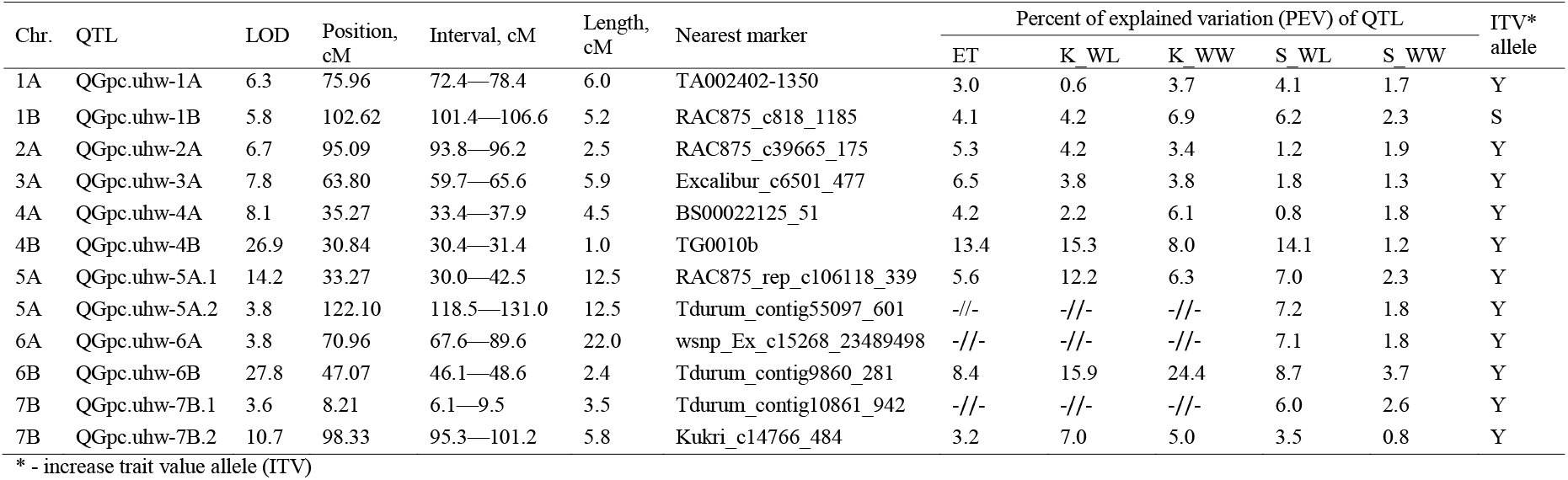
Summary of the QTLs associated with GPC in S×Y RIL population across five environments.

| Chr. | QTL | LOD | Position,<br>cM | Interval, cM | Length,<br>cM | Nearest marker | Percent of explained variation (PEV) of QTL |  |  |  |  | ITV*<br>allele |
| --- | --- | --- | --- | --- | --- | --- | --- | --- | --- | --- | --- | --- |
|  |  |  |  |  |  |  | ET | K_WL | K_WW | S_WL | S_WW |  |
| 1A | QGpc.uhw-1A | 6.3 | 75.96 | 72.4—78.4 | 6.0 | TA002402-1350 | 3.0 | 0.6 | 3.7 | 4.1 | 1.7 | Y |
| 1B | QGpc.uhw-1B | 5.8 | 102.62 | 101.4—106.6 | 5.2 | RAC875_c818_1185 | 4.1 | 4.2 | 6.9 | 6.2 | 2.3 | S |
| 2A | QGpc.uhw-2A | 6.7 | 95.09 | 93.8—96.2 | 2.5 | RAC875_c39665_175 | 5.3 | 4.2 | 3.4 | 1.2 | 1.9 | Y |
| 3A | QGpc.uhw-3A | 7.8 | 63.80 | 59.7—65.6 | 5.9 | Excalibur_c6501_477 | 6.5 | 3.8 | 3.8 | 1.8 | 1.3 | Y |
| 4A | QGpc.uhw-4A | 8.1 | 35.27 | 33.4—37.9 | 4.5 | BS00022125_51 | 4.2 | 2.2 | 6.1 | 0.8 | 1.8 | Y |
| 4B | QGpc.uhw-4B | 26.9 | 30.84 | 30.4—31.4 | 1.0 | TG0010b | 13.4 | 15.3 | 8.0 | 14.1 | 1.2 | Y |
| 5A | QGpc.uhw-5A.1 | 14.2 | 33.27 | 30.0—42.5 | 12.5 | RAC875_rep_c106118_339 | 5.6 | 12.2 | 6.3 | 7.0 | 2.3 | Y |
| 5A | QGpc.uhw-5A.2 | 3.8 | 122.10 | 118.5—131.0 | 12.5 | Tdurum_contig55097_601 | -/- | -/- | -/- | 7.2 | 1.8 | Y |
| 6A | QGpc.uhw-6A | 3.8 | 70.96 | 67.6—89.6 | 22.0 | wsnp_Ex_c15268_23489498 | -/- | -/- | -/- | 7.1 | 1.8 | Y |
| 6B | QGpc.uhw-6B | 27.8 | 47.07 | 46.1—48.6 | 2.4 | Tdurum_contig9860_281 | 8.4 | 15.9 | 24.4 | 8.7 | 3.7 | Y |
| 7B | QGpc.uhw-7B.1 | 3.6 | 8.21 | 6.1—9.5 | 3.5 | Tdurum_contig10861_942 | -/- | -/- | -/- | 6.0 | 2.6 | Y |
| 7B | QGpc.uhw-7B.2 | 10.7 | 98.33 | 95.3—101.2 | 5.8 | Kukri_c14766_484 | 3.2 | 7.0 | 5.0 | 3.5 | 0.8 | Y |

\* - increase trait value allele (ITV)

### Candidate gene analysis

The anchoring of the genetically mapped SNP markers into the physical position of WEW Zavitan pseudomolecules (Table S1) enabled us to define the physical intervals for each QTL (Table 5) and reveal genes residing within each interval (Table S2). We have estimated the physical intervals of the detected QTLs (1.5 LOD support interval) based on the proportion between the physical and genetic positions of the SNPs and of QTLs. The physical intervals of QTLs were ranged from 1.39 to 154.42 Mbp and the number of genes within these intervals varied from 5 to 653 (Table 5, Table S2). Of special interest are genes that are involved in transport and metabolism of nitrogen as potential CGs (Table S3). We have detected a total of 53 such CGs for the 12 detected QTL intervals (Table S3). The 20 most promising CGs are presented in Table 6.

**Table 5.**
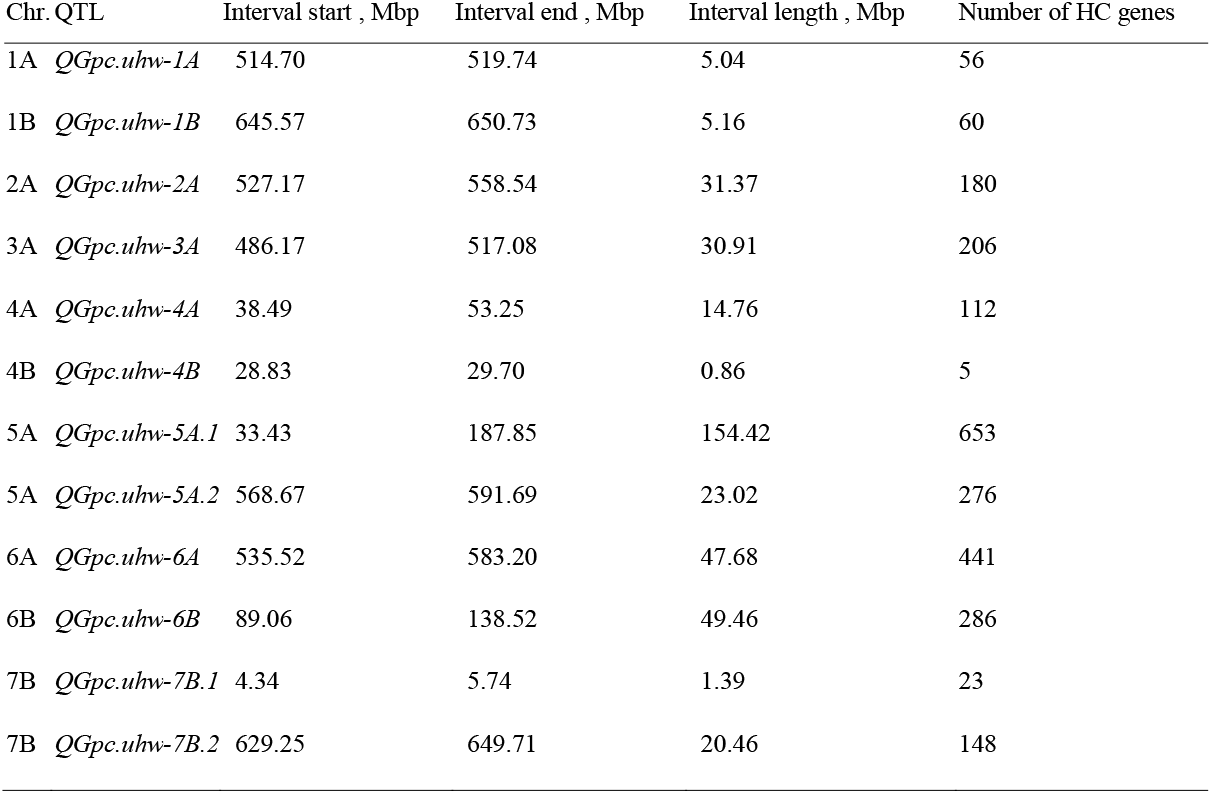
Physical intervals and number of genes within GPC QTLs in S×Y RIL population.

| Chr. QTL | Interval start , Mbp | Interval end , Mbp | Interval length , Mbp | Number of HC genes |
| --- | --- | --- | --- | --- |
| 1A *QGpc.uhw-1A* | 514.70 | 519.74 | 5.04 | 56 |
| 1B *QGpc.uhw-1B* | 645.57 | 650.73 | 5.16 | 60 |
| 2A *QGpc.uhw-2A* | 527.17 | 558.54 | 31.37 | 180 |
| 3A *QGpc.uhw-3A* | 486.17 | 517.08 | 30.91 | 206 |
| 4A *QGpc.uhw-4A* | 38.49 | 53.25 | 14.76 | 112 |
| 4B *QGpc.uhw-4B* | 28.83 | 29.70 | 0.86 | 5 |
| 5A *QGpc.uhw-5A.1* | 33.43 | 187.85 | 154.42 | 653 |
| 5A *QGpc.uhw-5A.2* | 568.67 | 591.69 | 23.02 | 276 |
| 6A *QGpc.uhw-6A* | 535.52 | 583.20 | 47.68 | 441 |
| 6B *QGpc.uhw-6B* | 89.06 | 138.52 | 49.46 | 286 |
| 7B *QGpc.uhw-7B.1* | 4.34 | 5.74 | 1.39 | 23 |
| 7B *QGpc.uhw-7B.2* | 629.25 | 649.71 | 20.46 | 148 |

**Table 6.** Information about selected CG that reside within the intervals of the detected QTLs

| Chr. | QTLs | IDs of candidate genes | Annotated function |
| --- | --- | --- | --- |
| 1A | *QGpc.uhw-1A* | TRIDC1AG048050 | Sulfite reductase [NADPH] hemoprotein beta-component |
| 1B | *QGpc.uhw-1B* | TRIDC1BG066150<br>TRIDC1BG066600 | glutamate receptor 3.4<br>Bifunctional inhibitor/lipid-transfer protein/seed storage 2S albumin superfamily protein |
| 2A | *QGpc.uhw-2A* | TRIDC2AG045010<br>TRIDC2AG047900 | Protein NRT1/ PTR FAMILY 4.3<br>Protein NRT1/ PTR FAMILY 6.4 |
| 3A | *QGpc.uhw-3A* | TRIDC3AG040540 | Protein NRT1/ PTR FAMILY 4.3 |
| 4A | *QGpc.uhw-4A* | TRIDC4AG007090<br>TRIDC4AG007800 | Protein transport protein GOT1<br>Protein transport protein Sec61 subunit alpha |
| 4B | *QGpc.uhw-4B* | TRIDC4BG006760 | SCARECROW-like 21 |
| 5A | *QGpc.uhw-5A.1* | TRIDC5AG007510<br>TRIDC5AG008560<br>TRIDC5AG005810<br>TRIDC5AG008600 | Protein NRT1/ PTR FAMILY 8.3<br>Protein NRT1/ PTR FAMILY 4.6<br>Protein NRT1/ PTR FAMILY 2.11<br>Protein NRT1/ PTR FAMILY 4.6 |
| 5A | *QGpc.uhw-5A.2* | TRIDC5AG056650 | Protein NRT1/ PTR FAMILY 4.5 |
| 6A | *QGpc.uhw-6A* | TRIDC6AG048910<br>TRIDC6AG049880 | nitrate reductase 1<br>nitrite reductase 1 |
| 6B | *QGpc.uhw-6B* | TRIDC6BG019590 | NAC domain protein (Gpc-B1) |
| 7B | *QGpc.uhw-7B.1* | TRIDC7BG000760<br>TRIDC7BG001030 | oligopeptide transporter 4<br>Nucleotide/sugar transporter family protein |
| 7B | *QGpc.uhw-7B.2* | TRIDC7BG057940<br>TRIDC7BG058370 | Protein NRT1/ PTR FAMILY 5.1<br>amino acid transporter 1 |

### Comparison of GPC QTLs from two *T. durum* x WEW RIL populations

We have compared the results of the GPC QTL analysis obtained in the current study for the S×Y population with our previous result obtained for another *T. durum* (Langdon) × WEW (G16-18) population (L×G) (Fatiukha et al. 2019a) in order to identify possible common QTLs based on co-location of QTL physical intervals. A total of 4 out of 12 QTLs identified in the current study (*QGpc.uhw-5A.1, QGpc.uhw-6A, QGpc.uhw-6B*, and *QGpc.uhw-7B.2*) were co-localized with QTLs identified in L×G population (Fig. 3). The major QTLs in both populations coincided with the position of *Gpc-B1* gene on chromosome 6B. Bearing in mind the allopolyploid nature of wheat genome we have search for homoeologous QTLs between populations using a list of homologous genes from Avni et al. (2017). We consider *QGpc.uhw-3A* and *QGpc.uhw-5A.1* in S×Y population as possible homoeologous for 3B.3 and 5B.2 QTLs in L×G population, respectively (Table S2). Interestingly, QTL *QGpc.uhw-5A.1* located on 5AS was colocalized with QTL 5A.2 and showed potential homoeology with QTL 5B.2 in L×G population.

**Figure 3.**
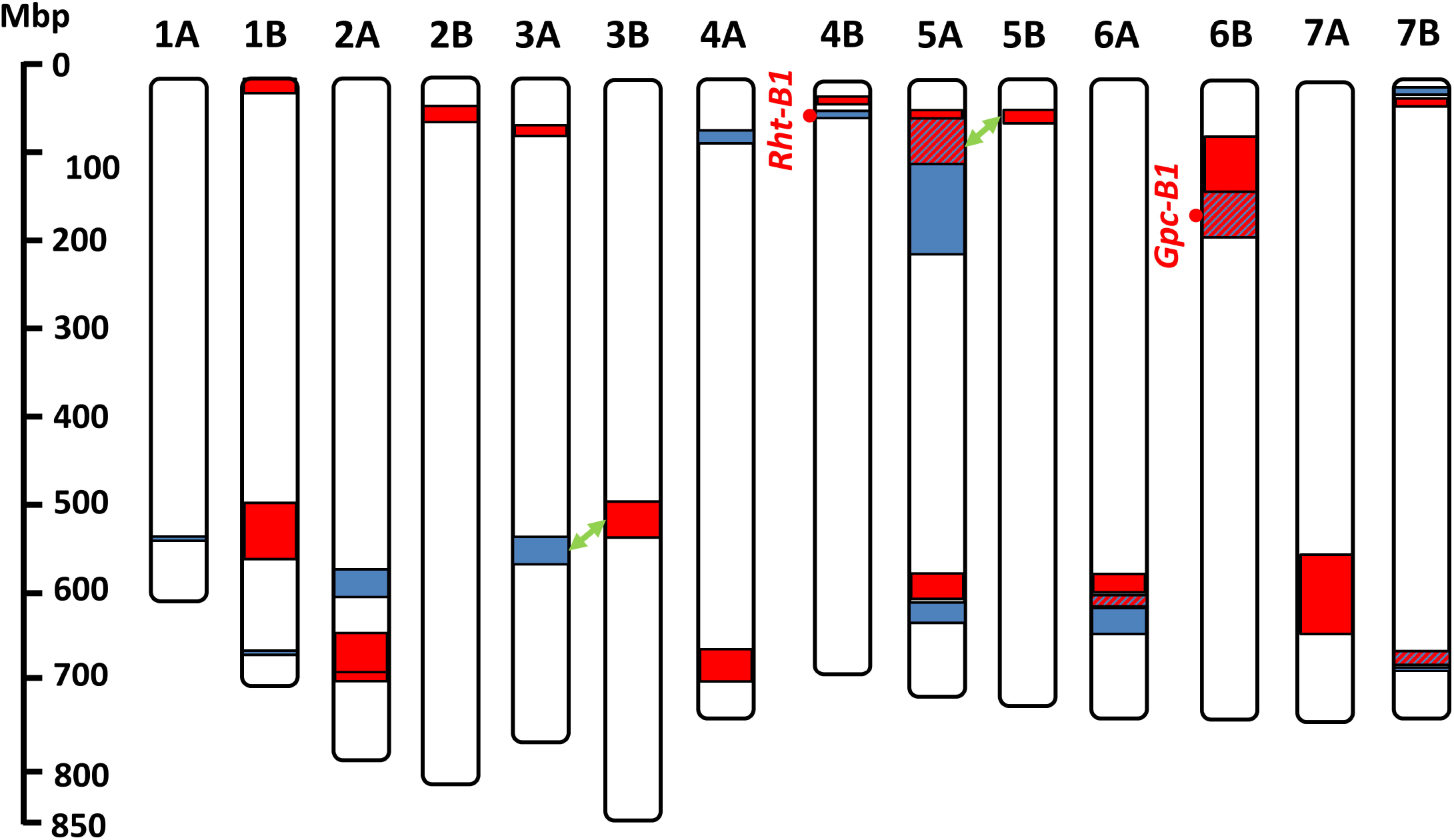
Physical intervals of GPC QTLs identified in S×Y (blue) and L×G (red) RIL populations. Co-localization of physical interval marked by diagonal lines; potential homoeologous QTLs are connected by green double arrow.

## Discussion

Increased protein content is one of the major objectives of grain quality improvement in wheat breeding. However, genetic improvement of wheat cultivars for many traits is limited due to genetic bottleneck (Peng et al. 2011), which is the result of early domestication events and subsequent selection in favor of yield related traits, thus, improving GPC requires broadening of wheat genetic diversity. This goal can be achieved by exploitation of wild crop relatives in breeding programs and genetic dissection of complex traits in order to upgrade food quality (Longin and Würschum 2016). The rich genepools of WEW (Krugman et al. 2018; Klymiuk et al. 2019b), emmer wheat *T. dicoccum* (Fedak 2015), and other wild relatives (Alvarez and Guzman 2018) were shown to be promising sources for genetic improvement of wheat. The detection of chromosomal regions responsible for increasing GPC is the first step towards this goal, followed by subsequent introgression of favorable alleles, cloning of promising genes, and revealing the underlying molecular mechanisms. The current study presents the construction of a high-density genetic map and QTL analysis of GPC for *T. durum*×WEW RIL population that demonstrated a high potential of WEW genepool for improvement of wheat quality.

Recent advantages in the development of high-throughput SNP genotyping in wheat (Wang et al. 2014; Winfield et al. 2016) led to a considerable improvement in construction of consensus maps (Wang et al. 2014; Maccaferri et al. 2015). Furthermore, the availability of reference genome assemblies of tetraploid (Avni et al. 2017) and hexaploid wheat (Appels et al. 2018) made it possible to compare the genetic and physical positions of markers and identify the genes residing within the intervals of interest. The high collinearity of the S×Y genetic map, constructed in the current study, with wheat consensus maps, as well as with the assembly of WEW pseudomolecules, highlights the quality and reliability of this map. Furthermore, genotyping of two RIL populations, derived from crosses of *T. durum* x WEW, using the 15K SNP array, resulted in comparable numbers of skeletal markers: 1,369 for L×G (Fatiukha et al. 2019a) and 1,510 for S×Y populations. The total length of the presented genetic map for S×Y (2,168 cM) is slightly longer than the map constructed for the L×G population (1,836 cM), and approximately equal to the length of the genetic map of the S×Z RIL population (2,111 cM), which is another *T. durum* x WEW cross, genotyped using the 90K SNP array (Avni et al. 2014a). Recent development in bioinformatics pipelines allows simple conversion of mapped SNPs to competitive allele-specific PCR (KASP) markers (Uauy et al. 2015) that can be used as flanking or even functional molecular markers (Klymiuk et al. 2019b), for marker-assisted breeding or fine mapping of the target QTLs.

GPC QTLs were reported previously for each chromosome of tetraploid and hexaploid wheat (Quraishi et al. 2017). Similarly, we detected GPC QTLs in 10 out of 14 chromosomes of tetraploid wheat. Our results showed clear advantage of WEW alleles for increasing GPC, compared to the corresponding durum allele, although Svevo is showing above average GPC for elite durum cultivars (Blanco et al. 2012). It is important to note that although WEW genepool was shown to be a valuable source for improving wheat GPC (Chatzav et al. 2010), only a few studies regarding genetic mapping of GPC in WEW were published. Among them, the identification of *Gpc-B1* QTL (Joppa et al. 1997) represents a great example for the exploitation of WEW genepool for GPC improvement, followed by the cloning of the NAC transcription factor, *NAM-B1*, underlying this QTL (Uauy et al. 2006). QTL analysis of the L×G RIL population revealed 8 loci with WEW alleles that conferred increased GPC, using low-density SSR-based map (Peleg et al. 2009). Further analysis, using high resolution SNP-based map, expanded the number of loci with contribution of WEW alleles from 8 to 14 (Fatiukha et al. 2019b). The identification of common or meta-QTLs between independent populations can help to improve QTL interval confidence, as demonstrated by Quraishi et al. (2017), and can be used for confirmation of detected effects in different genetic backgrounds. Nevertheless, the identification of co-located QTLs among the large number of publications is restricted by the absence of consensus positions for many of the published markers. In the current study, we were able to compare the physical QTL intervals, and the CGs that reside there, between the L×G and the S×Y GPC QTL maps, since the genetic markers were efficiently anchored to the reference Zavitan assembly of WEW. We found four possible common and two possible homoeologous QTLs between these two populations, as well as seven QTLs from S×Y population that were novel compared to L×G population. Since these two populations were developed using distant WEW genotypes, as well as different durum cultivars (Avni et al. 2014a), it is possible that the detected WEW alleles are absent or rare in cultivated wheat gene pool. Interestingly, the major QTLs identified in both populations were located on chromosome arm 6BS and were co-localized with the putative position of *Gpc-B1*. While the detected co-locations can be used as confirmation of QTL effects in different genetic backgrounds, such QTLs can still represent different alleles or even different CGs, conferring higher potential of broadening the GPC allele repertoire available for wheat improvement by exploitation of WEW genepool.

Many potential genes can be associated with GPC due to the complexity of the trait, and therefore, the proposed CGs should be treated carefully in further studies. Nevertheless, some of the CGs that showed strong functional association with GPC can be used as a basis for fine mapping, cloning and allele mining analysis. QTL analysis in RIL populations cannot provide single-gene resolution, however, we have identified two QTLs (*QGpc.uhw-4B* and *QGpc.uhw-7B.1*) with physical intervals around 1Mbp and only 5 and 23 genes residing within these intervals, respectively. Only one QTL interval in our study spanned over 154 Mbp that contain 653 genes, thus making it highly challenging to search for CG within this extremely large chromosome segment. The *Gpc-B1*, a known gene for increasing GPC in wheat (Tabbita et al. 2017), was found within the interval of a major GPC QTL (*QGpc.uhw-6B*) in the S×Y population. This gene is responsible for more efficient remobilization of nutrients from the senescing leaves to the grains during filling stage, thus, increasing protein and mineral accumulation in wheat grains (Uauy et al. 2006). Although, four *Gpc-B1* homoeologous and paralogous (Gpc-A1, Gpc-B1, Gpc-A2, and Gpc-B2 on chr. 6A, 6B, 2A, and 2B respectively) genes were identified in tetraploid wheat (Uauy et al. 2006) and showed effect on nutrient remobilization (Avni et al. 2014b), we did not detect copies of this gene within other QTL intervals in the S×Y population. Interestingly, a major dwarfing gene *Rht-B1* (Pearce et al. 2011) was identified within the shortest QTL interval with only five CGs. Taking into account that the cultivated parental line Svevo is a semi-dwarf cultivar carrying the dwarf allele of this gene (De Santis et al. 2018) and Y12-3 is a tall WEW (Peleg et al. 2005), we assume that this gene may exhibit a strong effect on all agronomical traits. Moreover, other studies in wheat also showed a possible effect of this gene on GPC, which makes *Rht-B1* a strong CG for the detected QTL (Fowler et al. 2016; Zou et al. 2017). Members of NRT1/PTR family of proton-coupled transporters, which are involved in nitrogen metabolism in different plant species (Fang et al. 2013; Naz et al. 2017; Castro-Rodriguez et al. 2017; Corratge-Faillie and Lacombe 2017) and, hence, influencing protein accumulation, were identified within five QTL intervals in the S×Y population and were selected as promising CGs. Sulfur metabolism was shown to be an important factor that can alter nitrogen accumulation (Howarth et al. 2008; Dai et al. 2015; Bonnot et al. 2017), following this idea glutamate and glutathione related CGs were identified within intervals of seven QTLs in the S×Y population and also was shown for QTLs in the L×G population (Fatiukha et al. 2019b).

### Conclusions and future perspectives

In the present study, we reported a QTL analysis study of GPC, based on a high-density SNP-based genetic map for the S×Y (durum×WEW) RIL population. We detected novel GPC QTLs that were not previously reported for durum×WEW populations and identified potential CGs for each of the 12 detected QTLs. Physical intervals of GPC QTLs in two durum×WEW populations were used for identification of four common and two homoeologous QTLs between them. *Gpc-B1* was characterized as a CG for a major GPC QTL confirming that this gene serves as one of the major sources of variation in GPC for wheat. The obtained results will serve as a basis for future introgression of favorable WEW alleles into bread and durum wheat, fine mapping, and cloning of promising QTLs.

Our results showed an advantage of WEW alleles for increasing GPC that emphasize the high potential of WEW genepool for improvement of modern wheat quality.

## Author contribution statement

A.F., A.B.K., C.P. T.F., and T.K. designed the research; I.L., G.L. performed field experiment and sample processing; A.F. performed the data analysis; A.F., V.K., T.K., and T.F wrote the manuscript.

## Acknowledgements

The research leading to these results has received funding from the European Community’s Seventh Framework Programme (FP7/ 2007-2013) under the grant agreement N°FP7-613556, Whealbi project; Carmel LTD and Kaiima Bio-Agritech Ltd, the Israeli Science Foundation (2289/16), and the Israeli field crops organization. We greatly acknowledge N. Filler, R. Jing-Jun and O. Chernjavska for their excellent technical assistance.

## Supporting Material

**Table S1.** Genetic and physical positions of the mapped SNPs

**Table S2.** List of High Confidence genes within QTL intervals and their homoeologs

**Table S3.** Summary of CGs

## References

Alvarez JB, Guzman C (2018) Interspecific and intergeneric hybridization as a source of variation for wheat grain quality improvement. Theor Appl Genet 131:225–251. doi: 10.1007/s00122-017-3042-x

Appels R, Eversole K, Feuillet C, et al (2018) Shifting the limits in wheat research and breeding using a fully annotated reference genome. Science 361:eaar7191. doi: 10.1126/science.aar7191

Asseng S, Ewert F, Martre P, et al (2014) Rising temperatures reduce global wheat production. Nat Clim Chang 5:143

Avni R, Nave M, Barad O, et al (2017) Wild emmer genome architecture and diversity elucidate wheat evolution and domestication. Science 357:93–97. doi: 10.1126/science.aan0032

Avni R, Nave M, Eilam T, et al (2014a) Ultra-dense genetic map of durum wheat × wild emmer wheat developed using the 90K iSelect SNP genotyping assay. Mol Breed 34:1549–1562. doi: 10.1007/s11032-014-0176-2

Avni R, Zhao R, Pearce S, et al (2014b) Functional characterization of GPC-1 genes in hexaploid wheat. Planta 239:313–324. doi: 10.1007/s00425-013-1977-y

Balla K, Rakszegi M, Li Z, et al (2011) Quality of Winter Wheat in Relation to Heat and Drought Shock after Anthesis. Czech J. Food Sci. 29: 117–128

Blanco A, Mangini G, Giancaspro A, et al (2012) Relationships between grain protein content and grain yield components through quantitative trait locus analyses in a recombinant inbred line population derived from two elite durum wheat cultivars. Mol Breed 30:79–92. doi: 10.1007/s11032-011-9600-z

Bonnot T, Bancel E, Alvarez D, et al (2017) Grain subproteome responses to nitrogen and sulfur supply in diploid wheat Triticum monococcum ssp. monococcum. Plant J 91:894–910. doi: 10.1111/tpj.13615

Carlos Brevis J, Dubcovsky J (2010) Effects of the Chromosome Region Including the Gpc-B1 Locus on Wheat Grain and Protein. Crop Sci 50:93–10

Castro-Rodriguez V, Canas RA, de la Torre FN, et al (2017) Molecular fundamentals of nitrogen uptake and transport in trees. J Exp Bot 68:2489–2500. doi: 10.1093/jxb/erx037

Chatzav M, Peleg Z, Ozturk L, et al (2010) Genetic diversity for grain nutrients in wild emmer wheat: potential for wheat improvement. Ann Bot 105:1211–1220. doi: 10.1093/aob/mcq024

Corratge-Faillie C, Lacombe B (2017) Substrate (un)specificity of Arabidopsis NRT1/PTR FAMILY (NPF) proteins. J Exp Bot 68:3107–3113. doi: 10.1093/jxb/erw499

Dai Z, Plessis A, Vincent J, et al (2015) Transcriptional and metabolic alternations rebalance wheat grain storage protein accumulation under variable nitrogen and sulfur supply. Plant J 83:326–343. doi: 10.1111/tpj.12881

De Santis M, Giuliani M, Giuzio L, et al (2018) Assessment of grain protein composition in old and modern Italian durum wheat genotypes. Italian Journal of Agronomy 13: 40–43

Doyle J (1991) DNA Protocols for Plants BT - Molecular Techniques in Taxonomy. In: Hewitt GM, Johnston AWB, Young JPW (eds). Springer Berlin Heidelberg, Berlin, Heidelberg, pp 283–293

Fang Z, Xia K, Yang X, et al (2013) Altered expression of the PTR/NRT1 homologue OsPTR9 affects nitrogen utilization efficiency, growth and grain yield in rice. Plant Biotechnol J 11:446–458. doi: 10.1111/pbi.12031

Fatiukha A, Deblieck M, Klymiuk V, et al (2019a) Genomic architecture of phenotypic plasticity of complex traits in tetraploid wheat in response to water stress. bioRxiv 565820. doi: 10.1101/565820

Fatiukha A, Klymiuk V, Peleg Z, et al (2019b) Variation in phosphorus and sulfur content shapes the genetic architecture and phenotypic associations within wheat grain ionome. bioRxiv 580423. doi: 10.1101/580423

Fedak G (2015) Alien Introgressions from wild Triticum species, T. monococcum, T. urartu, T. turgidum, T. dicoccum, T. dicoccoides, T. carthlicum, T. araraticum, T. timopheevii, and T. miguschovae. In: Molnár-Láng M, Ceoloni C, Doležel J (eds) Alien Introgression in Wheat: Cytogenetics, Molecular Biology, and Genomics. Springer International Publishing, Cham, pp 191–219

Flagella Z, Giuliani MM, Giuzio L, et al (2010) Influence of water deficit on durum wheat storage protein composition and technological quality. Eur J Agron 33:197–207. doi: https://doi.org/10.1016/j.eja.2010.05.006

Fowler DB, N’Diaye A, Laudencia-Chingcuanco D, Pozniak CJ (2016) Quantitative Trait Loci Associated with Phenological Development, Low-Temperature Tolerance, Grain Quality, and Agronomic Characters in Wheat (Triticum aestivum L.). PLoS One 11:e0152185

Golan G, Hendel E, Méndez Espitia GE, et al (2018) Activation of seminal root primordia during wheat domestication reveals underlying mechanisms of plant resilience. Plant Cell Environ 41:755–766. doi: 10.1111/pce.13138

Henry RJ, Nevo E (2014) Exploring natural selection to guide breeding for agriculture. Plant Biotechnol J 12:655–662. doi: 10.1111/pbi.12215

Howarth JR, Parmar S, Jones J, et al (2008) Co-ordinated expression of amino acid metabolism in response to N and S deficiency during wheat grain filling. J Exp Bot 59:3675–3689. doi: 10.1093/jxb/ern218

Huang L, Raats D, Sela H, et al (2016) Evolution and Adaptation of Wild Emmer Wheat Populations to Biotic and Abiotic Stresses. Annu Rev Phytopathol 54:279–301. doi: 10.1146/annurev-phyto-080614-120254

Joppa LR, Du C, Hart GE, Hareland GA (1997) Mapping Gene(s) for Grain Protein in Tetraploid Wheat (Triticum turgidum L.) Using a Population of Recombinant Inbred Chromosome Lines. Crop Sci 37:1586–1589. doi: 10.2135/cropsci1997.0011183X003700050030x

Jorgensen C, Luo M-C, Ramasamy R, et al (2017) A High-Density Genetic Map of Wild Emmer Wheat from the Karaca Dağ Region Provides New Evidence on the Structure and Evolution of Wheat Chromosomes. Front Plant Sci 8:1798. doi: 10.3389/fpls.2017.01798

Kao C-H, B Zeng Z, D Teasdale R (1999) Multiple Interval Mapping for Quantitative Trait Loci. Genetics 152:1203–16

Klymiuk V, Fatiukha A, Fahima T (2019a) Wheat tandem kinases provide insights on disease resistance gene flow and host-parasite co-evolution. Plant J:. doi: 10.1111/tpj.14264

Klymiuk V, Fatiukha A, Huang L, et al (2019b) Durum Wheat as a Bridge Between Wild Emmer Wheat Genetic Resources and Bread Wheat. In: Miedaner T, Korzun VBT-A of G and GR in C (eds) Woodhead Publishing Series in Food Science, Technology and Nutrition. Woodhead Publishing, pp 201–230

Korol A, Mester D, Frenkel Z, Ronin Y (2009) Methods for Genetic Analysis in the Triticeae. In: Feuillet C, Muehlbauer GJ, editors. Genetics and genomics of the Triticeae. New York: Springer Science + Business Media. pp . pp. 163–199

Krugman T, Chague V, Peleg Z, et al (2010) Multilevel regulation and signalling processes associated with adaptation to terminal drought in wild emmer wheat. Funct Integr Genomics 10:167–186. doi: 10.1007/s10142-010-0166-3

Krugman T, Nevo E, Beharav A, et al (2018) The Institute of Evolution Wild Cereal Gene Bank at the University of Haifa. Isr J Plant Sci 65:129–146. doi: https://doi.org/10.1163/22238980-00001065

Krugman T, Peleg Z, Quansah L, et al (2011) Alteration in expression of hormone-related genes in wild emmer wheat roots associated with drought adaptation mechanisms. Funct Integr Genomics 11:565–583. doi: 10.1007/s10142-011-0231-6

Longin CFH, Würschum T (2016) Back to the Future – Tapping into Ancient Grains for Food Diversity. Trends Plant Sci 21:731–737. doi: https://doi.org/10.1016/j.tplants.2016.05.005

Maccaferri M, Ricci A, Salvi S, et al (2015) A high-density, SNP-based consensus map of tetraploid wheat as a bridge to integrate durum and bread wheat genomics and breeding. Plant Biotechnol J 13:648–663. doi: 10.1111/pbi.12288

Mester D, Ronin Y, Minkov D, et al (2003) Constructing large-scale genetic maps using an evolutionary strategy algorithm. Genetics 165:2269–2282

Muqaddasi QH, Brassac J, Börner A, et al (2017) Genetic Architecture of Anther Extrusion in Spring and Winter Wheat. Front Plant Sci 8:754. doi: 10.3389/fpls.2017.00754

Nave M, Avni R, Ben-Zvi B, et al (2016) QTLs for uniform grain dimensions and germination selected during wheat domestication are co-located on chromosome 4B. Theor Appl Genet 129:1303–1315. doi: 10.1007/s00122-016-2704-4

Naz M, Fan X, Fan X, et al (2017) Plant nitrate transporters: from gene function to application. J Exp Bot 68:2463–2475. doi: 10.1093/jxb/erx011

OECD/FAO (2018) OECD-FAO Agricultural Outlook 2018-2027 - Special Focus: Middle East and North Africa

Pearce S, Saville R, Vaughan SP, et al (2011) Molecular characterization of Rht-1 dwarfing genes in hexaploid wheat. Plant Physiol 157:1820–1831. doi: 10.1104/pp.111.183657

Peleg Z, Cakmak I, Ozturk L, et al (2009) Quantitative trait loci conferring grain mineral nutrient concentrations in durum wheat x wild emmer wheat RIL population. Theor Appl Genet 119:353–369. doi: 10.1007/s00122-009-1044-z

Peleg Z, Fahima T, Abbo S, et al (2005) Genetic diversity for drought resistance in wild emmer wheat and its ecogeographical associations. Plant Cell Environ 28:176–191. doi: 10.1111/j.1365-3040.2005.01259.x

Peleg Z, Saranga Y, Yazici A, et al (2008) Grain zinc, iron and protein concentrations and zinc-efficiency in wild emmer wheat under contrasting irrigation regimes. Plant Soil 306:57–67. doi: 10.1007/s11104-007-9417-z

Peng JH, Sun D, Nevo E (2011) Domestication evolution, genetics and genomics in wheat. Mol Breed 28:281. doi: 10.1007/s11032-011-9608-4

Quraishi UM, Pont C, Ain Q, et al (2017) Combined Genomic and Genetic Data Integration of Major Agronomical Traits in Bread Wheat (Triticum aestivum L.). Front Plant Sci 8:1843. doi: 10.3389/fpls.2017.01843

Ronin YI, Mester DI, Minkov DG, et al (2017) Building Ultra-High-Density Linkage Maps Based on Efficient Filtering of Trustable Markers. Genetics 206:1285–1295. doi: 10.1534/genetics.116.197491

Shewry PR, Hey SJ (2015) The contribution of wheat to human diet and health. Food energy Secur 4:178–202. doi: 10.1002/fes3.64

Tabbita F, Pearce S, Barneix AJ (2017) Breeding for increased grain protein and micronutrient content in wheat: Ten years of the GPC-B1 gene. J Cereal Sci 73:183–191. doi: https://doi.org/10.1016/j.jcs.2017.01.003

Triboï□Blondel A, Triboï E, Martre P (2003) Environmentally □ induced changes in protein composition in developing grains of wheat are related to changes in total protein content. J Exp Bot 54:1731–1742. doi: 10.1093/jxb/erg183

Uauy C, Caccamo M, Ramirez-Gonzalez RH (2015) PolyMarker: A fast polyploid primer design pipeline. Bioinformatics 31:2038–2039. doi: 10.1093/bioinformatics/btv069

Uauy C, Distelfeld A, Fahima T, et al (2006) A NAC Gene regulating senescence improves grain protein, zinc, and iron content in wheat. Science 314:1298–1301. doi: 10.1126/science.1133649

Wang S, Wong D, Forrest K, et al (2014) Characterization of polyploid wheat genomic diversity using a high-density 90,000 single nucleotide polymorphism array. Plant Biotechnol J 12:787–796. doi: 10.1111/pbi.12183

Winfield MO, Allen AM, Burridge AJ, et al (2016) High-density SNP genotyping array for hexaploid wheat and its secondary and tertiary gene pool. Plant Biotechnol J 14:1195–1206. doi: 10.1111/pbi.12485

Zou J, Semagn K, Iqbal M, et al (2017) QTLs associated with agronomic traits in the Attila × CDC Go spring wheat population evaluated under conventional management. PLoS One 12:e0171528

